# The impact of recent population history on the deleterious mutation load in humans and close evolutionary relatives

**DOI:** 10.1101/073635

**Authors:** Yuval B. Simons, Guy Sella

**Affiliations:** Department of Biological Sciences, Columbia University, New York, NY, USA

## Abstract

Over the past decade, there has been both great interest and confusion about whether recent demographic events—notably the Out-of-Africa-bottleneck and recent population growth—have led to differences in mutation load among human populations. The confusion can be traced to the use of different summary statistics to measure load, which lead to apparently conflicting results. We argue, however, that when statistics more directly related to load are used, the results of different studies and data sets consistently reveal little or no difference in the load of non-synonymous mutations among human populations. Theory helps to understand why no such differences are seen, as well as to predict in what settings they are to be expected. In particular, as predicted by modeling, there is evidence for changes in the load of recessive loss of function mutations in founder and inbred human populations. Also as predicted, eastern subspecies of gorilla, Neanderthals and Denisovans, who are thought to have undergone reductions in population sizes that exceed the human Out-of-Africa bottleneck in duration and severity, show evidence for increased load of non-synonymous mutations (relative to western subspecies of gorillas and modern humans, respectively). A coherent picture is thus starting to emerge about the effects of demographic history on the mutation load in populations of humans and close evolutionary relatives.

## Introduction

The recent demographic history of human populations is reflected in their distributions of genetic variation. For instance, Europeans and Asians harbor a greater fraction of high frequency variants compared to Africans, likely due to an ancient “Out-of-Africa” bottleneck [1–4], and all of these populations harbor numerous rare variants resulting from more recent explosive growth [4–9]. Genetic variation in human populations has also been affected by founder events [10–12], by inbreeding [13,14], and by extensive admixture among populations [15,16] and with archaic humans [17,18].

It is therefore natural to ask whether recent demographic events also affected the burden of deleterious mutations, leading it to differ among extant human populations. In addition to the observation that overall patterns of genetic diversity vary among populations with different demographic histories, theory suggests that, at equilibrium, the efficiency with which purifying selection removes deleterious variation is profoundly affected by population size and degree of inbreeding [19–23]. With data now available to address the question, the possibility that there exist differences in the burden of deleterious mutation among human populations has garnered considerable attention.

Answers to this question have been confusing, because many studies appear to reach conflicting conclusions (cf. [24–26]). In trying to sort out the source of the conflicts, we start by reviewing what is meant by the burden of deleterious mutations. Traditionally, this burden has been quantified in terms of the mutation load (sometime abbreviated by load below), defined as the proportional reduction in average fitness due to deleterious mutations [19,23,27–29]. Under a simple model that assumes one locus with fitnesses 1, 1-hs and 1-s, the load is

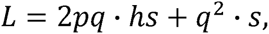

where *p* and *q* are the “normal” and “deleterious” allele frequencies. This reduces to

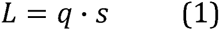

in the additive (semi-dominant) case and to

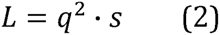

in the recessive case. More generally, load takes the form

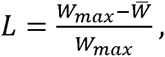

where *W*_*max*_ is the fitness of a mutation-free individual and 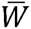 is the average fitness.

In practice, we cannot measure fitness and we know little about the distribution of selection and dominance coefficients, let alone about how the effects of deleterious mutations combine across loci. This is why recent studies have relied on population genetic summaries meant as proxies for load.

The choice of proxy turns out to be key in what the studies found. Notably, Lohmueller and colleagues (2008) introduced two summaries in order to compare individuals of European and African descent [30]. Using the first, they found the ratio of non-synonymous to synonymous segregating sites to be greater in the European sample than in the African one, which they interpreted as evidence for a reduced “efficacy of selection” in Europeans. Using the second summary, they found that European individuals carry (on average) more sites that are homozygous for non-synonymous derived alleles than do Africans, which they took as suggesting that Europeans likely suffer from a greater burden of recessive deleterious non-synonymous mutations. More recently, Simons et al. [31] and Do et al. [32] introduced a third summary (defined slightly differently in the two studies), the average number of derived non-synonymous variants per individual. They found no significant differences between European and African populations, and interpreted the pattern as indicating little or no difference in load. These studies and others [4,10,11, 33–42] applied the same methodologies to subsets of non-synonymous variants classified according to their predicted severity (e.g., using computational tools that rely on phylogenetic conservation and protein structure [43]), as well as to other human populations (see also [44,45]). With few exceptions (see below), analyses relying on the Lohmueller et al. summaries found substantial differences among populations whereas those that relied on the Simons et al. and Do et al. summaries did not.

## Comparing proxies for load

Given that the answer seems to depend on the summary, the question becomes which summary is most appropriate. The ideal summary would relate to load as directly as possible but also be insensitive to other differences among populations.

With these criteria in mind, we first consider the ratio of the number of non-synonymous to synonymous sites segregating in a population sample, *P*_*N*_/*P*_*S*_ (or subsets of non-synonymous sites). The idea is that *P*_*S*_ measures neutral diversity levels, and therefore *P*_*N*_/*P*_*S*_ measures an effective proportion of neutral non-synonymous mutations [46]; increased *P*_*N*_/*P*_*S*_ then reflects relaxed selection on non-synonymous mutations [25,30]. This interpretation applies at demographic equilibrium, e.g., when the population size is constant, or when non-synonymous mutations are either neutral or strongly selected, but it breaks down when neither assumption holds, which is precisely the case of interest. Under a population bottleneck, for example, *P*_*N*_/*P*_*S*_ first exhibit drastic changes because increased drift affects *P*_*N*_ and *P*_*S*_ differently, due to the different initial nonsynonymous and synonymous frequency spectra [30,32]; then *P*_*N*_ and *P*_*S*_ approach equilibrium at different rates because selected alleles have faster turnover than neutral ones [26,31,47,48]. Neither of these effects is related to relaxation of selection or increased load. To complicate the interpretation of *P*_*N*_/*P*_*S*_ even further, this statistic is extremely sensitive to the sample size (again because of the different synonymous and nonsynonymous frequency spectra). As a result, changes to PN/PS do not correspond to changes to load in any obvious way, even under straightforward demographic scenarios and assumptions about selection (Fig. 1A and [24]).

**Figure 1.**
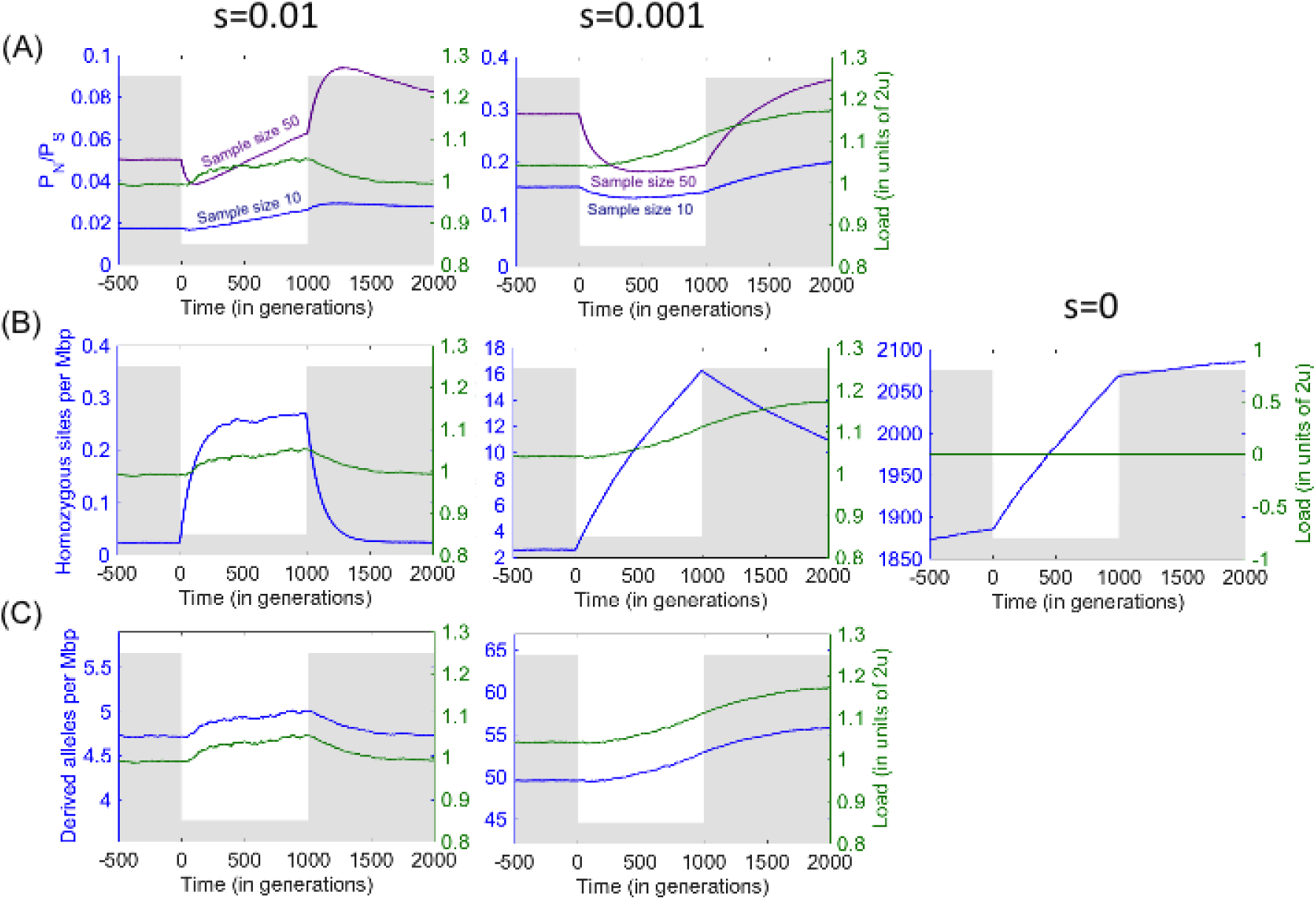
Load and summaries used to measure load under simulations with a bottleneck and additive selection. The population size (grey) drops from 10,000 to 1,000 at time 0 and recovers a 1,000 generations later. The number of sites simulated in each case was chosen to achieve standard errors below 1%. For further simulation details see [31].

We next consider the behavior of the average number of homozygous derived non-synonymous sites. For recessive deleterious mutations, individual load is related directly to this number (cf. Eq. 2). However, not all mutations that contribute to this summary are recessive or deleterious. Notably, many derived non-synonymous alleles may be neutral, and because they reach higher frequencies than deleterious ones, they would contribute disproportionally to the number of homozygous, derived sites, swamping any underlying signal. Moreover, demographic events often have marked effects on the number of neutral derived homozygous sites (Fig. 1B). For instance, bottlenecks increase the variance of neutral allele frequencies, *V*(*p*^*2*^)=*E*(*p*^*2*^)-*E*^*2*^(*p*), without affecting their mean, *E(p)*, thus increasing the frequency of homozygotes, *E*(*p*^*2*^). Comparing European populations that experienced the Out-of-Africa bottleneck to African ones that have not, we would therefore expect a large excess of homozygous, derived neutral sites in Europeans, even in the absence of a difference in load. Restricting the analysis to subsets of variants that are predicted to be more damaging will not solve this problem: while such subsets will include fewer neutral alleles, those neutral alleles that remain will contribute proportionally more homozygous sites, because more damaging variants have lower average frequencies [31]. Moreover, even if considering more damaging classes of variants helps to weed out neutral variants, non-recessive, deleterious variants continue to contribute, again complicating the relationship between the number of homozygous sites and load (Fig. 1B). Thus, the utility of this summary is severely compromised by its sensitivity to factors that have little to do with load.

Lastly, we consider the average number of non-synonymous derived alleles. This number is directly related to individual load when derived alleles are deleterious and additive (i.e., semi-dominant) (Fig. 1C). Moreover, for this summary, comparisons between populations are not confounded by the presence of neutral alleles [31,32]. This advantage becomes clear by considering a single sample from each population at a non-recombining locus (Fig. 2): if the mutation rate on the lineages leading from their common ancestor to both samples is the same then the expected number of neutral mutations on each lineage should also be the same. Using this statistic, no significant differences between human populations are seen for (putatively neutral) synonymous derived alleles (Fig. 3; [31,32] but see [49]). As we discuss below, non-synonymous variation is likely to be dominated by additive (or at least approximately additive) and neutral mutations, for which this summary is particularly well-suited.

**Figure 2.**
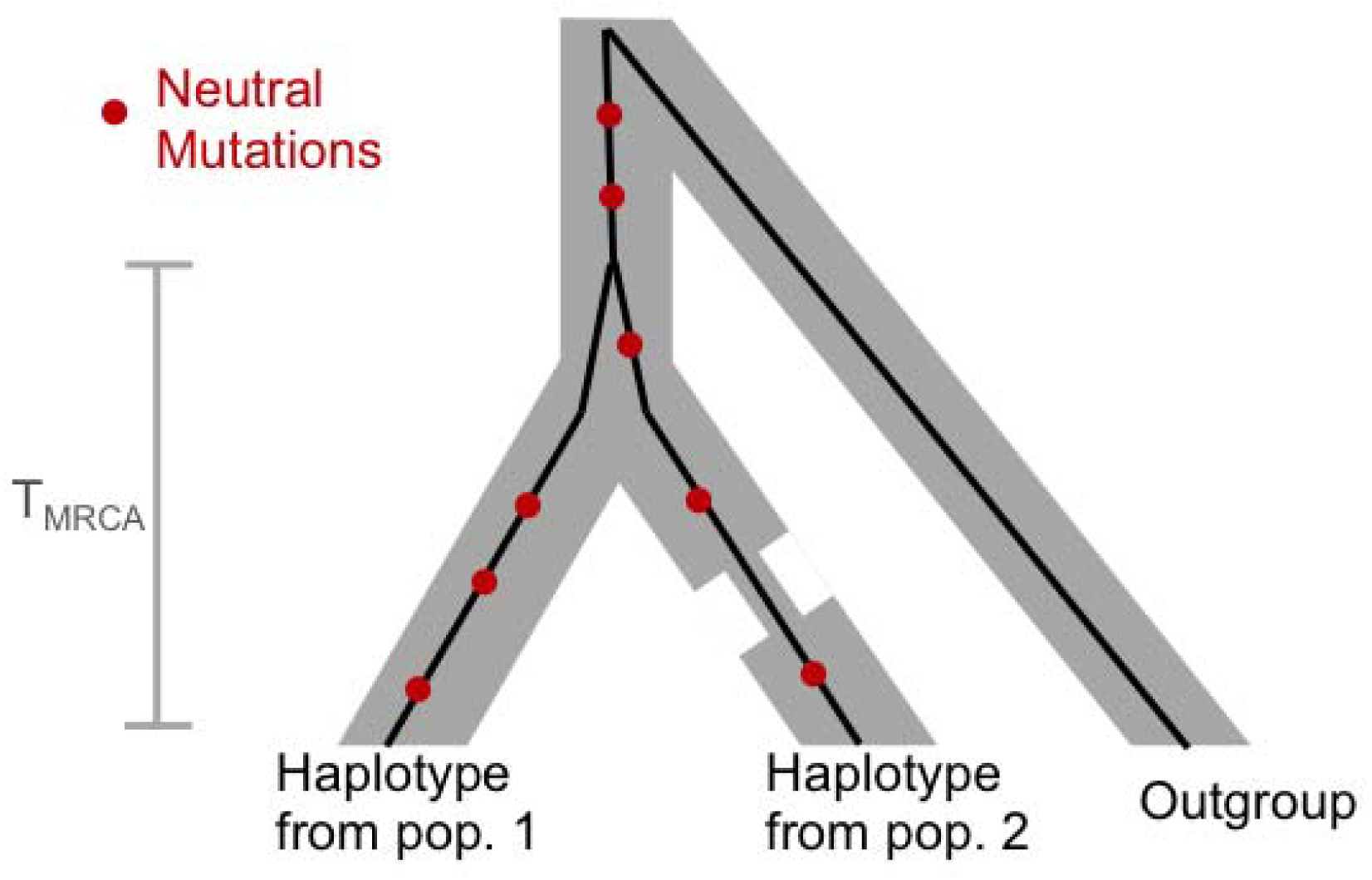
The genealogy at a locus of one sample from each of two populations, illustrating that the expected number of derived neutral alleles on each sample is the same, and depends only on the time to the most recent common ancestor (T_MRCA_) of the two lineages and not on the demographic history of the populations.

**Figure 3.**
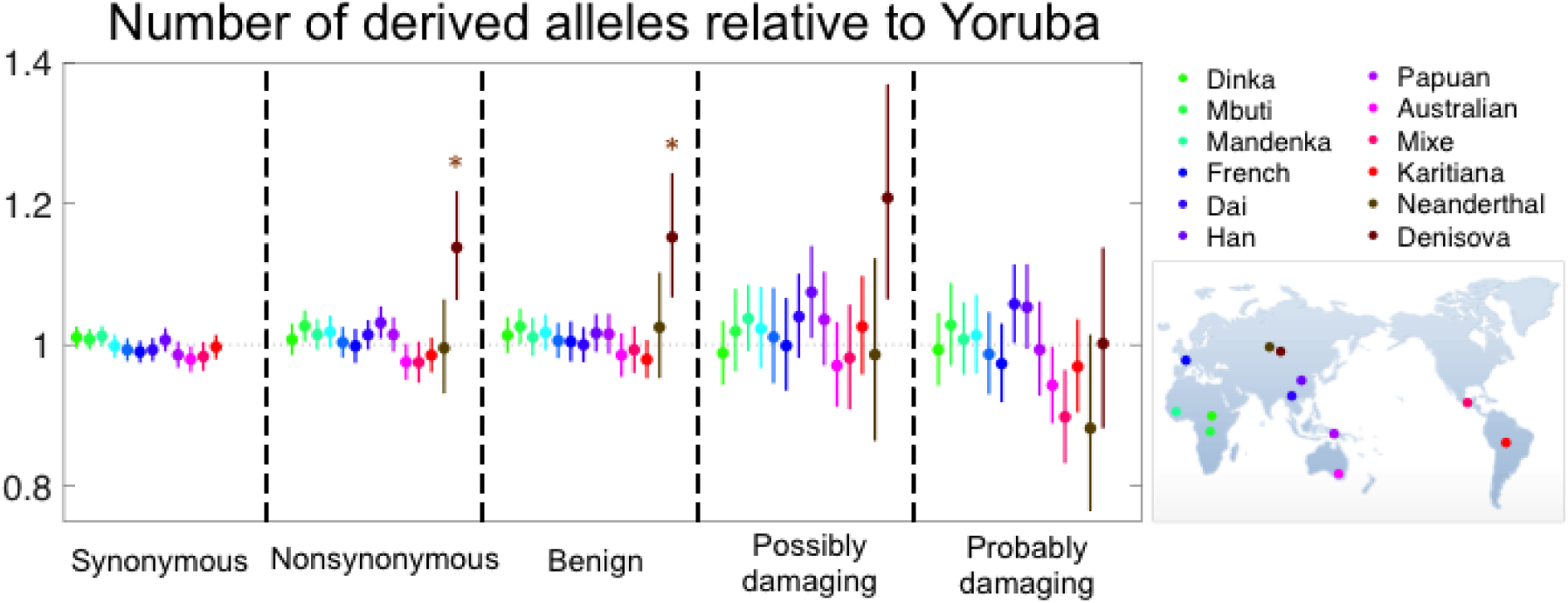
No significant difference between human populations in the mean number of derived alleles per individual. Each population sample was compared with the Yoruba sample, using data from [32]. Nonsynonymous SNP_s_ were classified into benign, possibly and probably damaging using PolyPhen 2.0 as described in [32]. The numbers of derived alleles for each comparison were counted at sites that were segregating in the joint sample from the two populations, and significance and the shown 95% confidence intervals were evaluated as described in Box 1, dividing the genome into 1,000 blocks. While there are multiple tests performed, it is not obvious how to correct for them, because population samples are also not independent. However, if we assume that at minimum 6 tests were performed then none of the comparisons among human populations is significant at the 5% level. The comparisons with Neanderthal and Denisova incorporate the modifications described in [32], where Denisova are significantly different regardless of the correction for multiple testing.

These considerations suggest that among summaries used to date, the number of derived non-synonymous alleles is a much more appropriate measure of load. Importantly then, studies that use this summary (or equivalent ones) found no significant differences among human populations (see Box 1), even though they analyzed large datasets from many populations and considered a variety of classifications of non-synonymous variants (Fig. 3). In the two exceptions of which we are aware, one study performed no statistical test of significance [40] and the other used a test that we believe does not fully account for uncertainty [36] (see Box 1). Of course, other factors could potentially affect this summary (recessivity of deleterious mutations is considered below). However, in order for them to result in there being no difference among populations, they would have to almost perfectly cancel out with any true signal. We therefore conclude that there is currently no reliable evidence for differences in load among human populations driven by differences in demographic history, at least at non-synonymous sites.

Observing little or no differences in load among populations might seem at odds with theoretical predictions. Specifically, theory predicts that at demographic equilibrium, a considerable portion of deleterious alleles for which *2N*_*es*_ ≤ 1 will be fixed, leading to a much greater load in smaller populations [19,22,54]. Consistent with the reduced efficacy of selection in smaller populations, lineages that tended to have smaller effective population sizes over long evolutionary timescales (e.g., since the split between rodents and primates) show evidence for relaxed constraint at coding and regulatory regions [55,56]. One might therefore expect a substantial increase in load, due to the additive mutations that the Out-of-Africa bottleneck turned from strongly to weakly selected. In fact, the duration of the bottleneck was too short to have led to many deleterious fixations, and therefore the increase is predicted to be minor (Fig. 1) [31]. A similar argument applies to the effects of explosive growth, which is much too recent to impact load [24,31,57]. More generally, the presumed duration of the demographic events that differ among human populations are much shorter than the timescales required for weakly selected variation to equilibrate (roughly on the order of one over the mutation rate; cf. [31]), which explains why the differences expected at equilibrium are not seen in data.

In contrast to the additive case, in the fully recessive case, the mutation load and average number of deleterious alleles can change rapidly and potentially dramatically [31,32,58-61]. As an illustration, consider variation that is strongly selected throughout a bottleneck and subsequent expansion (Fig. 4B). When the population size drops, the increase in genetic drift leads to a loss of variation and a consequent drop in the average number of deleterious alleles per individual. However, some deleterious alleles drift to higher frequencies, contributing disproportionally to the number of homozygous sites and causing a surge in load. The response to changing population size is faster when selection is stronger, with a new equilibrium approached on the timescale of allelic turnover 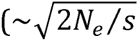 generations [31,60]). The load at equilibrium is insensitive to the population size, but the average number of deleterious alleles per individual scales with 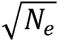. The expansion after the bottleneck is also accompanied by rapid changes to load and to the number of deleterious alleles (Fig. 4B).

**Figure 4.**
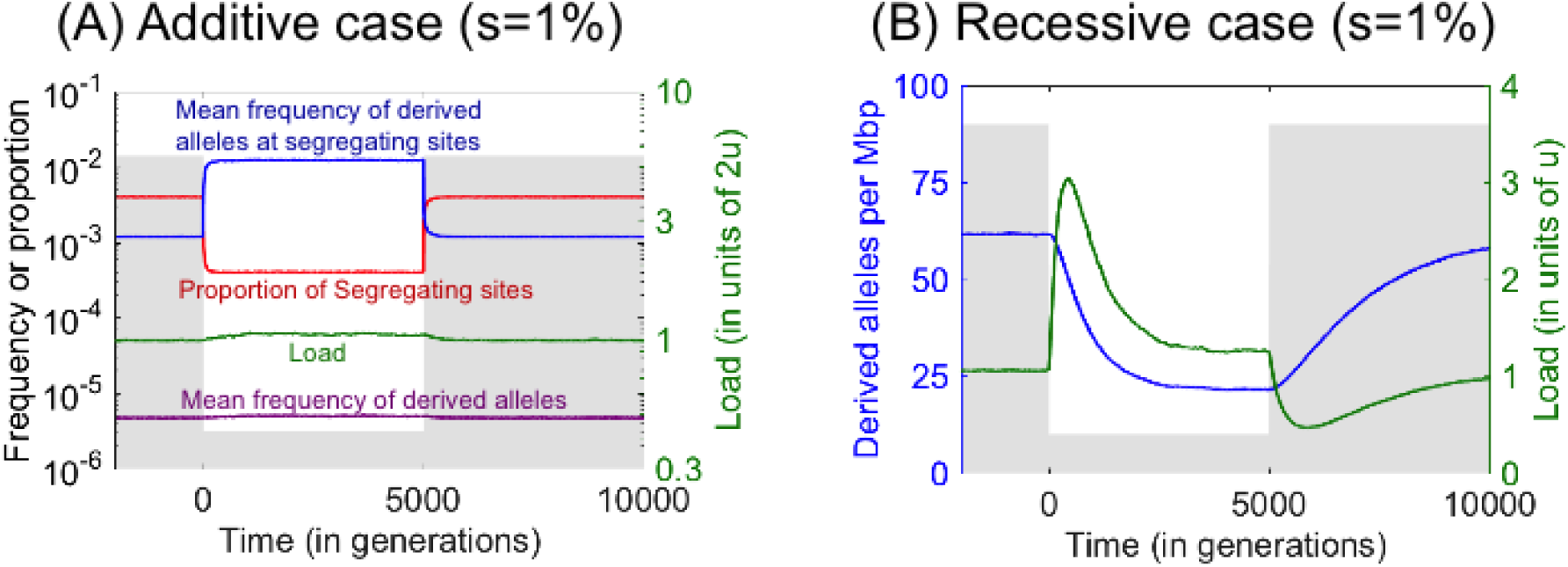
Bottleneck effects on load and the number of derived alleles in additive (A) and recessive (B) cases. The population size (grey) drops from 10,000 to 1,000 at time 0 and recovers 5,000 generations later. The number of sites simulated in each case was chosen to achieve standard errors below 1%. For further simulation details see [31].

#### Box 1: Statistical tests for demographically induced differences in load

Having chosen the number of derived alleles (of a given class) per individual as our proxy for load, the next question becomes how to test for significant differences between populations. One option is to test for a significant difference between population samples in the average counts per individual (e.g., using a Mann-Whitney test). The problem with this approach is that even if we were to run the exact evolutionary experiment twice, with the same demographic history, we would expect our summary to take different values, due to stochasticity in the mutational and genealogical processes. Thus, it is unclear whether a significant result reflects a difference in the demographic history of the two populations. An alternative approach is to divide the genomes into blocks, and bootstrap over these blocks to test for differences between the populations. The idea here is that different blocks represent independent evolutionary experiments with the same demographic history. The blocks need to be sufficiently large for different blocks to represent independent genealogies. They also need to be resampled at random from the union of the two population samples to control for the shared part of the genealogical history of the two populations and to ensure that other factors that vary along the genome (e.g., gene density) are the same in the population samples compared. It turns out that the two approaches to assessing statistical significance yield different answers: when the former approach is applied to sufficiently large samples, it indicates significant difference between populations [36], whereas the latter approach, which we argue is more appropriate, shows none.

## Should we expect substantial differences in load among human populations?

This question has been tackled using a combination of simulations and analytical tools. Most studies focused on changes to population size and specifically on bottlenecks and recent growth (but see [40]), with parameters based on rapidly improving demographic inferences [4,50–53]. Because much less is known about the distribution of selection and dominance coefficients, let alone about how the effects of deleterious mutations combine across loci, most studies considered single locus models under a range of selection and dominance coefficients (but see [25,36,53]). Even in this case, a general understanding of demographic effects on load requires reviewing many cases, notably different selection regimes (cf. [31]). Here we focus on some general insights that have emerged.

One important conclusion is that while bottlenecks and growth can dramatically affect both the number and frequencies of deleterious (and neutral) mutations, these shifts roughly cancel each other out, resulting in much subtler effects on load (Fig. 4A). In particular, for mutations that are effectively neutral (*2N*_*s*_<<1) or partially dominant and strongly selected (*2N*_*hs*_>>1) throughout the period being considered, changes to population size have been shown to have no effect on load [31]. Thus, it is quite possible to see little or no differences in load among populations, despite clear differences in overall patterns of genetic variation.

## Observing changes in recessive load

Given that bottlenecks and expansions occurred in the recent history of many human populations (e.g., [2,10–12]), why do we not see significant differences among populations in the number of derived non-synonymous alleles due to recessive variants? One possibility is that the proportion of recessive variants among derived alleles is too small for them to generate significant differences among populations. If this were the case, then we might hope to see such a difference when applying the test to subsets that are enriched for recessive deleterious variants.

Population genetics theory and experiments suggest that alleles with large effects are more likely to be recessive, whereas alleles with smaller effect are likely to be additive [62–67]. While non-synonymous variation is likely dominated by mutations of smaller effects, Loss of Function (LoF) mutations are plausibly enriched for mutations of large effects, and indeed there are numerous examples of recessive LoF Mendelian diseases [68,cf. 69]. In turn, as discussed above, theory predicts that strongly selected recessive variation responds to changes in population size fairly quickly. In accordance with theory, founder populations, such as Ashkenazi Jews and Finns, are known to have their own disease heritage, i.e., a particularly high incidence of specific recessive diseases, but fewer segregating recessive alleles underlying these and other recessive diseases than in neighboring outgroup populations ([12,70–75], cf. [69]). We may therefore expect to see differences in the average number of LoF alleles per individual between founder and outgroup populations.

Providing strong evidence to this effect in humans, Narasimhan et al. [14] found significantly fewer LoF alleles per individual in Finns than in other (non-founder) populations. They also found significantly fewer LoF mutations in the Birmingham/Bradford Pakistani heritage population than in other populations (except Finns), likely due to a recent surge in inbreeding (which, similar to a bottleneck followed by an expansion, increases homozygosity [76–78], leading to the more efficient purging of recessive deleterious mutations [79,80]). In gorillas, Xue et al. [81] inferred that eastern lowland and mountain subspecies experienced a more drastic reduction in population size than their western lowland and cross river counterparts, with recent population sizes in eastern subspecies so small that autozygosity levels exceed those found in the most inbred human populations. As expected from this demographic history, they also find evidence for markedly and significantly fewer LoF mutations per individual in the two eastern subspecies than in their western counterparts.

Observing a reduction in the number of LoF mutations per individual does not necessarily imply that the load is currently lower. For example, as illustrated in Fig. 4B, the relationship between the average number of alleles per individual and recessive load is also affected by the demographic history since the bottleneck. Nonetheless, such an observation likely indicates a past change in load and, in conjunction with other information, e.g., about changes to population size or levels of inbreeding, inferences about the current load may be feasible.

## Observing changes in additive load

While theory predicts that the increase to the load of additive mutations following a bottleneck would be extremely slow, it may still be detectable if the drop in population size was sufficiently long and severe. One example is provided by the increased reduction in population size in eastern compared to western gorillas, which was estimated to have been more severe and longer lasting than the Out-of-Africa bottleneck in humans [52,81]. Eastern gorilla subspecies also carry significantly more non-synonymous alleles per individual than their western counterparts, suggesting that the greater reduction in population size also led to an increase in load due to additive mutations [81].

Similarly, both Neanderthals and Denisovans are inferred to have had an extremely low population size since their main separation from humans, which was longer ago than the separation of gorilla subspecies [18,82]. These estimates suggest that they should have also incurred a substantial increase in load. In fact, their effective population sizes are among the lowest measured in any taxon [83], leading to speculation that a corresponding accumulation of deleterious mutations may have contributed to their demise [84]. To test the hypothesis that their load was higher, Do et al. [32] compared the number of derived non-synonymous alleles per individual between these archaic hominins and modern humans, using a modified summary aimed to control for the branch shortening and DNA degradation in ancient samples [32]. They found a markedly and significantly greater number in Denisovans, indicative of increased load, but not in Neanderthals (Fig. 3).

Strong evidence for increased load in Neanderthals comes from another source: the distribution of Neanderthal ancestry along the genomes of modern humans. Current estimates suggest that 2-4% of the genome of non-African individuals is the result of introgression from Neanderthals and that Neanderthal haplotypes collectively span ~30% of the genome [17,85,86]. This Neanderthal ancestry is, on average, markedly lower in regions of high gene density and low recombination, indicating that, with some notable exceptions (cf. [87]), selection has been acting to remove Neanderthal DNA [85,86,88,89]. A couple of recent studies have shown that these ancestry patterns are plausibly explained by selection against deleterious mutations from Neanderthals [53,84] (not found by Do et al. [32], possibly due to technical complications in the use of ancient samples). Using the ancestry patterns to infer the strength of selection against non-synonymous Neanderthal mutations, Juric et al. [84] found that it accords with the range that would have been effectively neutral in Neanderthals but effectively selected in humans. These studies further suggest, albeit speculatively, that some residual “admixture load” may remain in non-Africans today.

## Conclusion

The contradictory conclusions about the impact of recent demographic events on the mutation load in humans can be traced back to the use of different summary statistics and to how they are affected by factors other than load. We contend that of the summaries used thus far, the number of derived alleles per individual is both more sensitive and specific to load. Applying this summary and appropriate statistical tests to data yields findings that accord with theoretical expectations for modern human populations, archaic humans and subspecies of gorilla. Specifically, the findings follow from the plausible assumption that non-synonymous variation is dominated by mutations of small additive effects, whose contribution to load should changes slowly in response to changes in population size, whereas LoF mutations are likely enriched for large recessive effects, whose contribution to load should respond rapidly to changes in population size or surges of inbreeding. Of course, future advances, such as improvements in our classification of variants, may reveal differences in load that evade us at present. However, current findings suggest that differences among extant human populations are likely to be small.

## Acknowledgements.

We thank Molly Przeworski, Guy Amster, Jeff Ross-Ibarra, Aylwyn Scally and Jonathan Pritchard for comments on the manuscript. We owe special thanks to Ron Do, for his considerable help in producing Figure 2, and to Vagheesh Narasimhan, for answering many questions about his analyses. This work was funded by NIH grant GM115889 to GS.

